# EXERCISE-INDUCED BENEFITS ON GLUCOSE HANDLING IN A MODEL OF DIET-INDUCED OBESITY ARE REDUCED BY CONCURRENT NICOTINAMIDE MONONUCLEOTIDE

**DOI:** 10.1101/2020.06.22.164459

**Authors:** Josephine Yu, D. Ross Laybutt, Lynn-Jee Kim, Lake-Ee Quek, Lindsay E. Wu, Margaret J. Morris, Neil A. Youngson

## Abstract

**Objective:** Almost 40% of adults worldwide are classified as overweight or obese. Exercise is a beneficial intervention in obesity, partly due to increases in mitochondrial activity, with a potential role for the concomitant increase in nicotinamide adenine dinucleotide (NAD^+^). Recent studies have shown that increasing NAD^+^ levels through pharmacological supplementation with precursors such as nicotinamide mononucleotide (NMN) improved metabolic health in high fat diet (HFD) fed mice. We examined the combined effects of NMN and treadmill exercise on the metabolic dysregulation in HFD-induced obesity.

**Methods:** Five-week old female C57BL/6J mice were exposed to control diet or HFD. Mice fed HFD were treated with NMN in drinking water (400mg/kg; HNMN), treadmill exercise (HEx) or combined NMN and exercise (HNEx).

**Results:** Unexpectedly, NMN administration impaired several aspects of exercise-induced benefits in HFD mice, including glucose tolerance, glucose stimulated insulin secretion from islets and reduced hepatic triglyceride accumulation. Mechanistically, HNEx mice displayed increased antioxidant and reduced prooxidant gene expression in both islets and muscle, suggesting that altered redox status is associated with the loss of exercise-induced health benefits with NMN co-treatment.

**Conclusion:** Our data show that NMN treatment blocks the beneficial metabolic effects of exercise in a mouse model of diet-induced obesity in association with disturbances in redox metabolism.

**Highlights:**

- NMN dampened exercise-induced benefits on glucose handling in diet-induced obesity.
- NMN administration in exercise enhanced ratio of antioxidants to prooxidants.
- We suggest NMN administration may not be beneficial when NAD^+^ levels are replete.

## 1. INTRODUCTION

Obesity is a major health concern of the 21^st^ century. The World Health Organization (WHO) reported the worldwide incidence of obesity to be nearly tripled in 2016 compared to 1975, with over 1.9 billion adults classified as overweight, 650 million of whom were obese [1]. Obesity is a major risk factor for a multitude of other non-communicable diseases such as cardiovascular disease, non-alcoholic fatty liver disease, type 2 diabetes and osteoarthritis [2, 3]. Although multiple factors can contribute, the increased prevalence of obesity is thought to be primarily driven by an energy imbalance whereby more energy rich foods are being consumed but energy expenditure is reduced [1]. There is a plethora of evidence for the ability of exercise to overcome adverse effects of obesity, including improved glucose homeostasis, insulin sensitivity and liver fat accumulation [4–7]. Thus, exercise is currently a widely used intervention to aid weight loss and improve patient wellbeing.

In the context of obesity and high fat diet (HFD) feeding, the excessive consumption of nutrients is thought to drive mitochondrial dysfunction. One pathway which may be responsible for this is oxidative stress, by which an increase in reactive oxygen species (ROS) compromises mitochondrial function through reductions in mitochondrial biogenesis and impaired oxidation efficiency [8–10]. Mitochondria play a central role in oxidative metabolisms which involves the catabolism of fuel (lipids, proteins, and carbohydrates) to produce adenosine triphosphate (ATP), the main form of cellular energy. An integral molecule in this process is nicotinamide adenine dinucleotide (NAD^+^), a critical redox cofactor which has recently gained increased interest after the recognition of its role in metabolism and DNA repair [11]. It is thought that the detrimental health consequences of obesity and aging may in part be driven by a reduction in NAD^+^ levels, impairing fundamental metabolic reactions and the activity of NAD^+^ consuming enzymes such as the poly (ADP-ribose) polymerases (PARPs) and sirtuins (SIRTs), and in particular the mitochondrial sirtuin SIRT3 [12, 13]. In recent years, there has therefore been growing interest in boosting NAD^+^ levels to improve mitochondrial function.

Exercise benefits health in diet-induced obesity in both human and animal models. In humans, this is mediated through enhanced liver function and reduced insulin resistance [14, 15]. In rodent models, exercise benefits multiple systems including suppressed inflammatory cell recruitment in adipose tissue, prevention of diastolic dysfunction and restoration of insulin clearance [4, 16, 17]. One proposed mechanism is the ability of exercise to restore NAD^+^ levels and increase mitochondrial biogenesis which are typically lowered in obesity [18–20]. Exercise has been shown to increase NAD^+^ levels at multiple sites including liver, adipose tissue and skeletal muscle [21–23]. These changes were strongly associated with beneficial effects on metabolic outcomes including improved glucose tolerance. Rodent models have also shown that exercise training can increase expression of genes associated with mitochondrial biogenesis, such as upregulation of muscle *Pgc1*α [24].

The association between increased NAD^+^ levels and exercise-induced health benefits has resulted in pharmacological strategies to increase NAD^+^ levels through exogenous administration of NAD^+^ precursors such as nicotinamide mononucleotide (NMN) and nicotinamide riboside (NR). Many animal models have demonstrated beneficial effects of NMN administration in both ageing and diet-induced metabolic disease [13, 25]. In these mouse models, increased NAD^+^ was strongly associated with metabolic benefits such as improved glucose tolerance. Previous work by our group showed that administering NMN by intraperitoneal injection lessens the impact of diet-induced obesity, by improving glucose tolerance and reducing hepatic lipid accumulation [23]. NMN administration and exercise were independently able to increase NAD^+^ levels, and each intervention provided tissue-specific benefits. Thus, in this study, we aimed to examine the potential synergistic effects of exercise and chronic, oral administration of NMN in female mice challenged with HFD-induced obesity.

## 2. MATERIALS AND METHODS

### 2.1 Animal Experimentation

Female C57BL/6J mice were purchased from the Animal Resources Centre (Canning Vale, WA, Australia). All procedures were approved by the UNSW Animal Ethics Committee (ACEC no. 17/73B). Four-week old mice were allowed one week to acclimatize to their new environment before being allocated to either a standard chow (14kJ/g digestible energy, 12% from lipids; Gordon’s Specialty Feeds) or HFD (19kJ/g digestible energy, 43% from lipids; SF04-001 Specialty Feeds). Mice were housed four per cage in (temperature: 21±2°C; light/dark cycle: 12hr/12hr), with food and water available *ad libitum*. After 10 weeks of diet, when HFD-fed mice were more than 30% heavier than chow-fed mice, interventions were started. The HFD-fed mice were then split into four groups: HFD, HFD with NMN (HNMN), HFD with exercise (HEx) and HFD with both NMN and exercise (HNEx). Water consumption was measured prior to the start of the NMN intervention for HNMN and HNEx groups. Based on these calculations, in addition to the body weights which were measured weekly, NMN (GeneHarbor, Hong Kong) was provided in drinking water (400mg/kg body weight) from 15 weeks of age until 23 weeks of age. Water intake was monitored to confirm the dissolved NMN did not affect water intake, and to maintain the correct dosage as bodyweights changed.

Treadmill exercise started at 15 weeks of age for HEx and HNEx groups. Mice were allowed 4 days of acclimatization and training on treadmills (Exer3/6; Columbus Instruments) before starting the training regimen of exercise 6 days a week for 8 weeks. Mice ran for 50 minutes a day, starting with a 5-minute warm up period where the treadmill speed started at 6m/min and increased to 10m/min after 3 minutes. Mice then ran at 15m/min for 20 minutes and were allowed a 5-minute break whereby the treadmill was slowed to 6m/min, before resuming their run at 15m/min for another 20 minutes. Control, HFD and HNMN mice experienced time in a stationary treadmill 6 days a week.

### 2.2 Body Composition and Respiratory Quotient

At 17 weeks of age, each mouse underwent a whole-body composition analysis by nuclear magnetic resonance imaging (EchoMRI) assessing their fat mass and lean mass. At 18 weeks of age, respiratory quotient and resting energy expenditure were determined using comprehensive laboratory animal monitoring systems (CLAMS; Columbus Instruments) over a 24-hour period after acclimatization in the cages.

### 2.3 Glucose Tolerance Test (GTT)

GTT was performed at 20 weeks of age following 6 hours fasting. Tail vein blood was collected by tail nick and basal glucose levels were measured by glucometer (Accu-Chek Performa; Roche Diabetes Care). 2g/kg lean mass of glucose (30% glucose solution) was administered by intraperitoneal injection and blood glucose was monitored for 2 hours.

### 2.4 Tissue Collection and Islet Isolation

HEx and HNEx mice were exercised on treadmill the day before cull. Mice were fasted overnight before cull. Mice were anaesthetized with ketamine/xylazine mix (200/20 mg/kg i.p.), blood was collected by cardiac puncture before decapitation. Pancreatic islets were collected as previously described using liberase digestion (Roche) and gradient separation (Ficoll-Paque) before being handpicked under a microscope [26]. Groups of 5 freshly isolated islets were collected in quadruplicate for immediate use in a glucose stimulated insulin secretion assay. The remaining islets were collected for RNA extraction. All collected tissues were snap frozen in liquid N_2_ and stored at −80°C until use.

### 2.5 Glucose Stimulated Insulin Secretion (GSIS)

Isolated islets were incubated in Krebs Ringer Bicarbonate buffer (136mM NaCl, 4.7mM KCl,1.2mM KH_2_PO_4_, 5mM NaHCO_3_, 1.2mM MgSO_4_.7H_2_O, 1M HEPES and 1M CaCl_2_) containing 0.1% BSA for 1 hour at 37°C. Basal readings were taken, and islet cells were treated with KRB containing 2mM glucose followed by 20mM glucose. Each treatment consisted of a 1-hour incubation at 37°C. Collected secretions were assayed for insulin release using an insulin radioimmunoassay kit (RI-13K, Merck) as per manufacturer’s protocol.

### 2.6 Triglyceride Assay

Triglyceride content in the liver was measured as previously described [23]. Samples were homogenized in choloroform:methanol (2:1) using a Precellys 24 homogenizer. Samples were washed and centrifuged, the organic phase was evaporated under nitrogen then the extract re-dissolved in ethanol. Triglyceride content was measured using triglyceride reagent (Roche Diagnostics) against a glycerol standard curve (G7793-5ML, Sigma-Aldrich). Results were read at 495nm.

### 2.7 NAD Assay

Levels of NAD^+^ and NADH were quantified in liver as previously described [23]. Briefly, samples were homogenized, and an aliquot was taken for protein quantification. Samples were then extracted using phenol:chloroform:isoamyl alcohol and chloroform. An aliquot was taken to quantify total NAD, which was left on ice, and another to quantify NAD^+^ which was acidified with HCl, heated at 65°C then neutralized with NaOH. Samples were transferred to a 96-well plate, treated with alcohol dehydrogenase, then quantified using the iMark microplate reader. Protein was quantified as previously described [23] using standards (Thermo Scientific) and read at 595 nm.

### 2.8 Mitochondrial DNA (mtDNA) qPCR Assay

Powdered tissue samples were lysed (10mM Tris pH8, 100mM NaCl, 10mM EDTA, 0.5% SDS buffer) with 10mg/mL proteinase K overnight at 50°C. DNA was extracted as previously described [23]. Briefly, samples were mixed with phenol:chloroform. Supernatant was treated with 1μL of RNAse A at 37°C. Sodium acetate and absolute ethanol were added, and samples were incubated at −80°C, centrifuged and washed with 75% ethanol. Pellets were left to air dry then re-dissolved nuclease-free water. PCR was performed on the extracted DNA in duplicate, using *CytB* (F: CCCACCCCATATTAAACCCG; R: GAGGTATGAAGGAAAGGTATTAGGG) as a marker of the mitochondrial genome and Bioline SensiFAST SYBR No-Rox Mix (BIO-98020). A control sample was generated by mixing an aliquot of DNA from each sample and diluted to create a standard curve. PCR was performed on the Roche LightCycler 480 (Roche Diagnostics, Australia) and readings were normalized to a marker on the nuclear genome, *36b4* (F: ACTGGTCTAGGACCCGAGAAG; R: TCAATGGTGCCTCTGGAGATT).

### 2.9 Quantitative RT-PCR

Quadricep muscle was homogenized in 1mL of TRI-Reagent® (93289-25ML; Sigma-Aldrich) using a Precellys 24 tissue homogenizer (Bertin Instruments) as per manufacturer’s protocol. Islet RNA was extracted using Qiagen miRNeasy mini kits (217004) as per manufacturer’s protocol. RNA content was quantified using a DS-11 spectrophotometer (DeNovix Inc.).

One microgram (quadricep) or 0.2 micrograms (islets) of RNA was treated with DNAse I Amplification Grade (AMPD1-1KT; Sigma-Aldrich) before being reverse transcribed using the High Capacity cDNA reverse transcription kit (4368814, Applied Biosystems) as per manufacturer’s protocol. Samples were prepared in duplicate and expression of genes of interest were assessed using primers as listed: *Pgc-1a* (F: TATGGAGTGACATAGAGTGTGCT; R: CCACTTCAATCCACCCAGAAAG), *Ppara* (F: TACTGCCGTTTTCACAAGTGC; R: AGGTCGTGTTCACAGGTAAGA), *Sod2* (F: TTAACGCGCAGATCATGCA; R: GGTGGCGTTGAGATTGTTCA) and *Nox4* (F: GAAGATTTGCCTGGAAGAACC; R: AGGTTTGTTGCTCCTGATGC). A control sample was made by combining an aliquot of cDNA from each of the samples, then diluted to generate a standard curve for each of the target genes. Gene expression levels were quantified using the Roche Lightcycler 480. All readings were normalized to the geometric mean of two housekeeping genes for each sample - *Ywhaz* (F: TAGGTCATCGTGGAGGGTCG; R: GAAGCATTGGGGATCAAGAACTT), and *Hprt* (F: TCAGTCAACGGGGGACATAAA; R: GGGGCTGTACTGCTTAACCAG) for quadriceps; *Ywhaz* and *Tbp* (F: AGTGCATACTTTATCACCAAGCA; R: CCACTTGTACGAACTGGGTTTT) for islets. Housekeeping genes were selected from 5 possible genes (including *Gapdh* and *Rpl27*) using the Normfinder package for R (MOMA).

### 2.10 Thiobarbituric Acid Reactive Substances (TBARS)

Quadriceps muscle was homogenized in RIPA buffer using a Precellys 24 tissue homogenizer. Samples were nutated for 2 hours at 4°C then centrifuged at 10,000g for 10 minutes. Supernatant was extracted and protein concentration was measured as previously described. TBARS, by-products of lipid peroxidation, was assessed by measuring malondialdehyde (MDA) concentration, an end-product of lipid peroxidation, using the OxiSelect™ TBARS assay kit (Cell Biolabs Inc.) as per manufacturer’s protocol.

### 2.11 LC-MS/MS

Quadricep muscle was homogenized in cold extraction buffer (HPLC-grade acetonitrile-methanol-water mixture 2:2:1). Samples were centrifuged at 16,000g for 10 minutes at 4L and supernatants were dried under speed vacuum (Savant SpeedVac SPD140DDA Vacuum Concentrator, Thermo Fisher Scientific) before resuspension in LC-MS-grade water.

Supernatant was analyzed by LC-MS/MS where liquid chromatography was performed by HPLC (1260; Agilent) with XBridge™ Amide column (3.5 μm, 2.1 × 100 mm, Waters Corporation). LC separation was performed as previously reported [27] with minor modifications. For the mobile phase, the pH of Buffer A was adjusted to pH 5 using 20mM acetic acid instead of 20mM ammonium hydroxide to achieve better separation of acidic NAD^+^ metabolites from their amide counterparts. Buffer B was 100% acetonitrile with flow rate set at 200μl/min and flowed in gradient mode with percentages of Buffer B set at 85% (0 min), 85% (0.1 min), 70% (10 min), 30% (13 min), 30% (17 min), 85% (17.5 min), and 85% (30 min). The injection volume was 2.5μL. HPLC was coupled to the QTRAP® 5500 mass spectrometer controlled by Analyst^®^ Software (AB SCIEX, Toronto, Canada) for detection of metabolites in multiple reaction monitoring (MRM) mode with ion source set at 350°C and ionization voltage at 4500 volts with positive/negative polarity switching. . For data analysis, MSConvert (version 3.0.18165-fd93202f5) and in-house MATLAB scripts were used to integrate metabolite peaks and extract data. Integrated peak areas were normalized by dividing the peak area of the sample by the peak area of the internal standard for each metabolite, before being further divided by frozen tissue weight.

### 2.12 Statistical Analysis

Data were analyzed either by one-way or repeated measures ANOVA with Tukey’s post-hoc tests using SPSS Statistics 24 software (IBM). Symbols indicate significant effects between groups (**\*** compared to control, # compared to HFD, † compared to HNMN, ‡ compared to HEx, ^ compared to HNEx) with the number of symbols represent significance levels (one symbol, *p* < 0.05; two symbols, *p* < 0.01; three symbols, *p* < 0.001).

## 3. RESULTS

### 3.1 Effects of oral NMN administration and exercise on body weight, composition and physiological parameters

As expected, HFD feeding significantly increased body weight (**Figure 1A**) and at 15 weeks of age, HFD fed mice were 30% heavier than control mice (HFD: 28.48 ± 0.41g, control: 21.64 ± 0.34g). At this point, HFD fed mice were split into their respective intervention groups. At the end of the study, at 24 weeks of age (**Figure 1B**), there was a 44% increase in body weight when comparing HFD mice with no intervention (33.72 ± 0.79g) to control mice (23.35 ± 0.47g). HNMN (33.85 ± 1.08g) and HNEx (33.02 ± 0.88g) mice had comparable body weight to HFD mice. At this time, after 9 weeks of exercise, body weight of HEx mice was reduced by 12% (29.79 ± 1.12g) relative to HFD mice. This reduction in body weight was not reflected in fat mass taken as a percentage of body weight (**Figure S1A**), but rather in lean mass (**Figure S1B**). Close monitoring in a CLAMS metabolic cage system showed similar trends in food (**Figure S2C**) and water (**Figure S2D**) intake across groups, with no significant effect of diet or interventions. Ambulatory activity (**Figure S2B**) also showed minimal variation between groups. We observed an effect of diet on respiratory exchange ratio determined by indirect calorimetry (**Figure S2A**), confirming standard chow fed mice utilized carbohydrates as their primary source of energy whereas HFD-fed mice utilized lipids as their main substrate, with no effects of either oral NMN administration or exercise.

**Figure 1:**
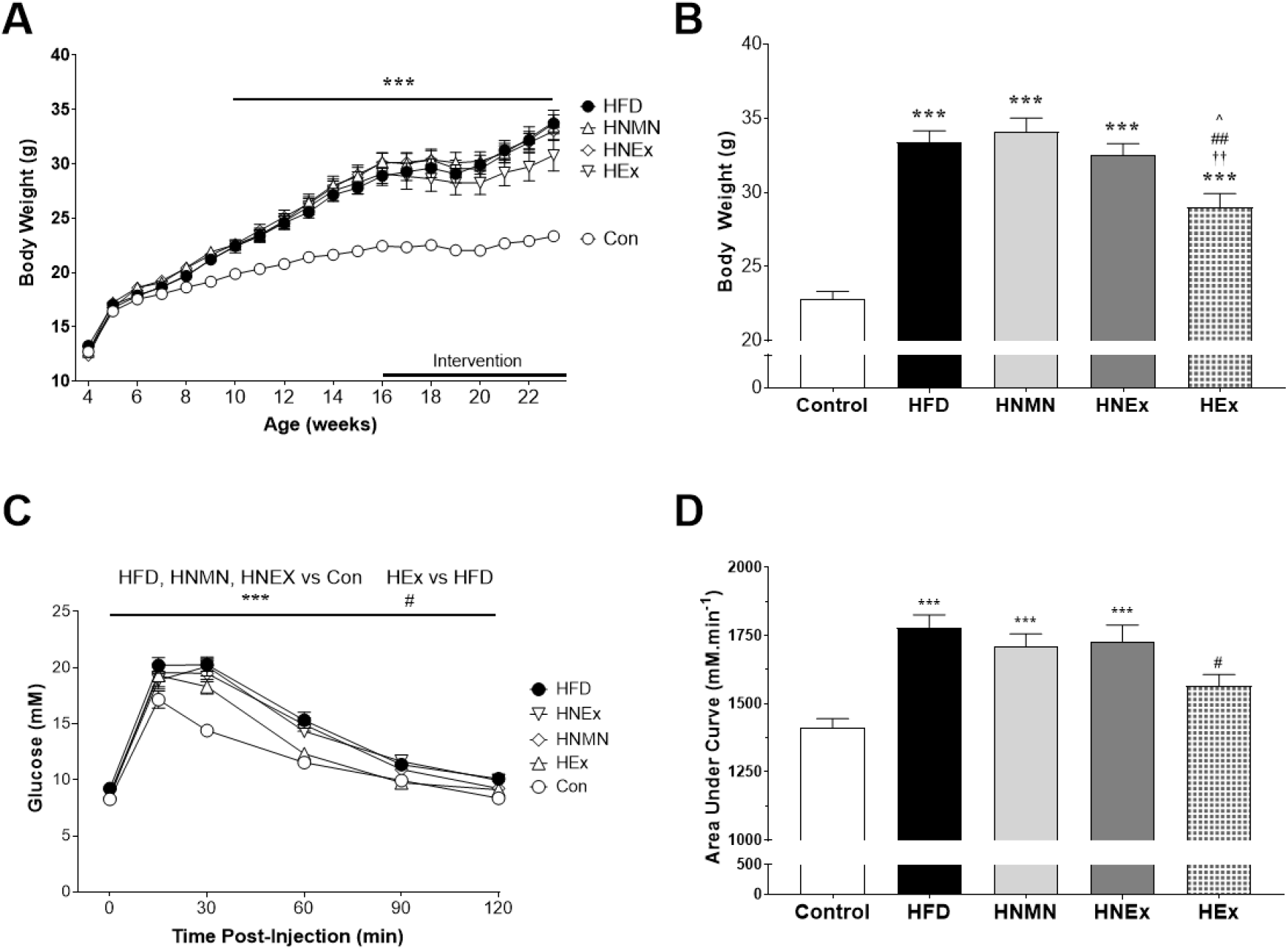
Effects of oral NMN administration and exercise on body weight (A and B), and glucose tolerance (C and D). Data expressed as mean ± SEM (n=14-16/group). Body weight over time (A) and glucose data collected over the duration of the GTT (C) were analyzed by repeated measures ANOVA followed by Tukey’s post-hoc test. Endpoint body weight at 24 weeks of age (B) and area under the curve (D) data were analyzed by one-way ANOVA followed by Tukey’s post-hoc test. * *p* < 0.05, *** *p* < 0.001 significant difference compared to control mice # *p* < 0.05, ## *p* < 0.01 significant difference compared to HFD mice †† *p* < 0.01 significant difference compared to HNMN mice ^ *p* < 0.05 significant difference compared to HNEx mice.

### 3.2 NMN co-treatment reduced the beneficial effects of exercise on HFD-induced glucose intolerance

No differences were detected in fasted basal glucose levels, regardless of diet. NMN administration alone did not have any observable benefits on HFD-induced glucose intolerance (**Figure 1C**), however treadmill exercise alone improved overall clearance (**Figure 1D**). Notably, this was abrogated in HNEx mice which received both exercise and oral NMN, suggesting that NMN abrogated the benefits of exercise.

### 3.3 Only exercise improved response of islets to glucose

We next examined the effects of exercise and NMN treatment alone and in combination on pancreatic beta cell function. Islets harvested from mice across groups secreted insulin in low glucose conditions (2 mM) at levels comparable to basal levels in chow control mice (**Figure 2A**). Challenging islets with 20 mM glucose (**Figure 2A**) resulted in a 5-fold increase in insulin secretion in chow fed animals, and as expected, this was decreased to a 2.5-fold increase in islets from HFD animals. Similar to the pattern observed in GTT, NMN administration alone was not able to improve GSIS, whereas exercise alone ameliorated the impairment caused by HFD, showing an improved response to 20 mM glucose (4-times chow basal). Interestingly, the beneficial effect of exercise on islet GSIS was negated in HNEx mice which again points towards NMN dampening the benefits of exercise.

**Figure 2:**
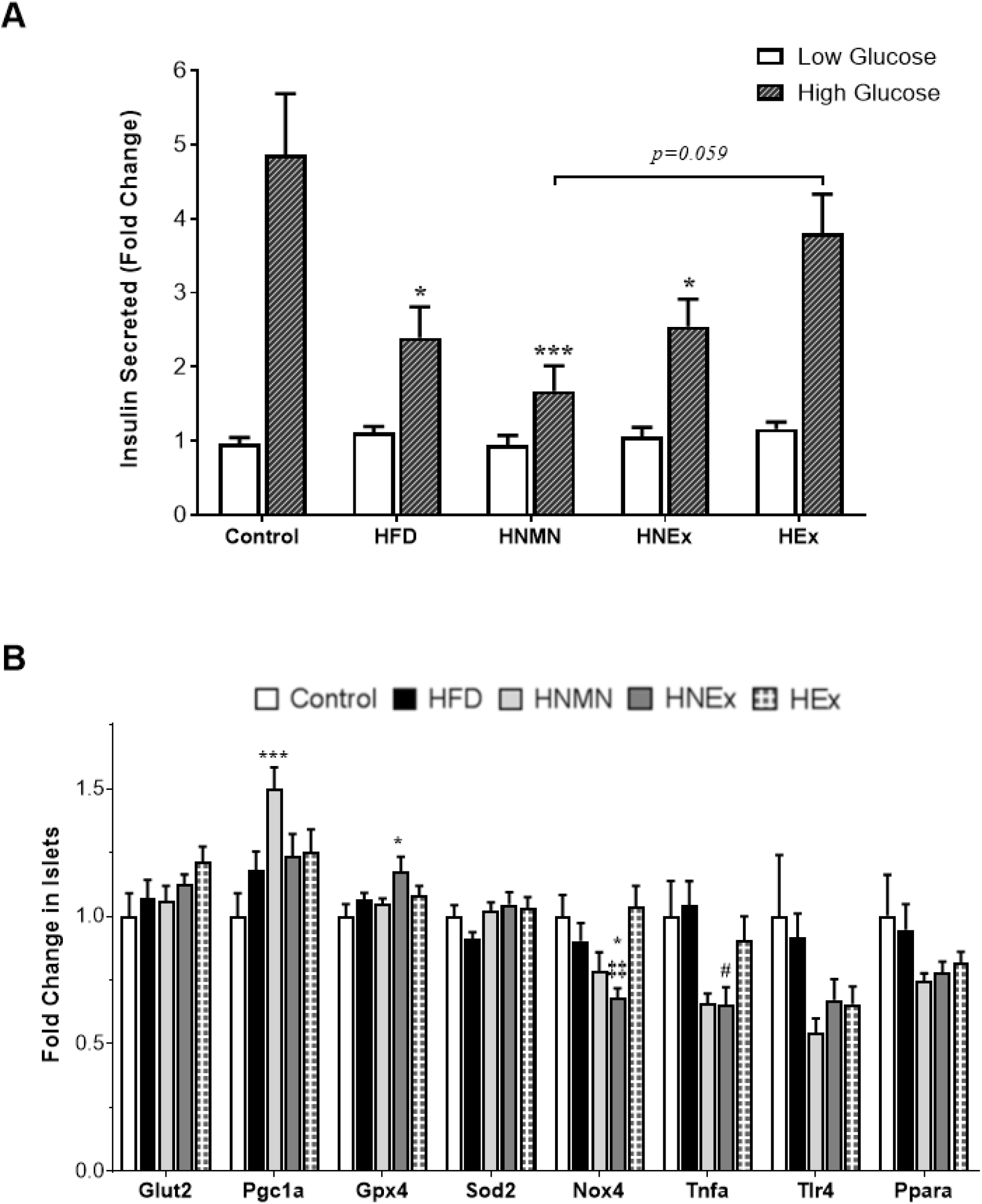
Effects of oral NMN administration and exercise on glucose-stimulated insulin secretion (A; n=10-12/group) and gene expression in islets (B; n=8-12/group). Data expressed as mean ± SEM in fold change compared to control mice. Data for each gene were normalized to two house-keeper genes and analyzed by one-way ANOVA followed by Tukey’s post-hoc test. *p* < 0.05, *** *p* < 0.001 significant difference compared to control mice # *p* < 0.05 significant difference compared to HFD mice ‡‡ *p* < 0.01 significant difference compared to HEx mice

We next sought to determine whether these changes were associated with altered expression of genes related to mitochondrial function, inflammation and ROS. Oral NMN administration at 400mg/kg significantly upregulated *Pgc1a* expression compared to chow fed mice (**Figure 2B**). We note that expression of *Tnfa* appeared to be downregulated under both NMN conditions. HNEx mice showed upregulated expression of the antioxidant enzyme glutathione peroxidase (*Gpx4*) and reduced expression of NADPH oxidase 4 (*Nox4*) compared to controls, suggesting an increased ratio of antioxidant to prooxidant gene expression. This differential regulation suggests a change in redox status that favors enhanced capacity for handling ROS, which can act as a signal for GSIS [28, 29]. Thus, our studies raise the possibility that disturbance of redox metabolism may be involved in NMN-associated dampening of the beneficial effects of exercise on GSIS.

### 3.4 Endpoint measures

Quadriceps mass (**Table 1**) was not different across groups, however it was observed to be significantly decreased in HFD fed groups when corrected for body weight, although exercise alone improved this. Mice fed HFD had over four times the retroperitoneal fat mass of control mice, with a slightly smaller increase in HEx mice (*p<0.001*, **Table 1**). These differences remained when fat mass was corrected for overall body weight (*p<0.001*, **Table 1**).

**Table 1:**
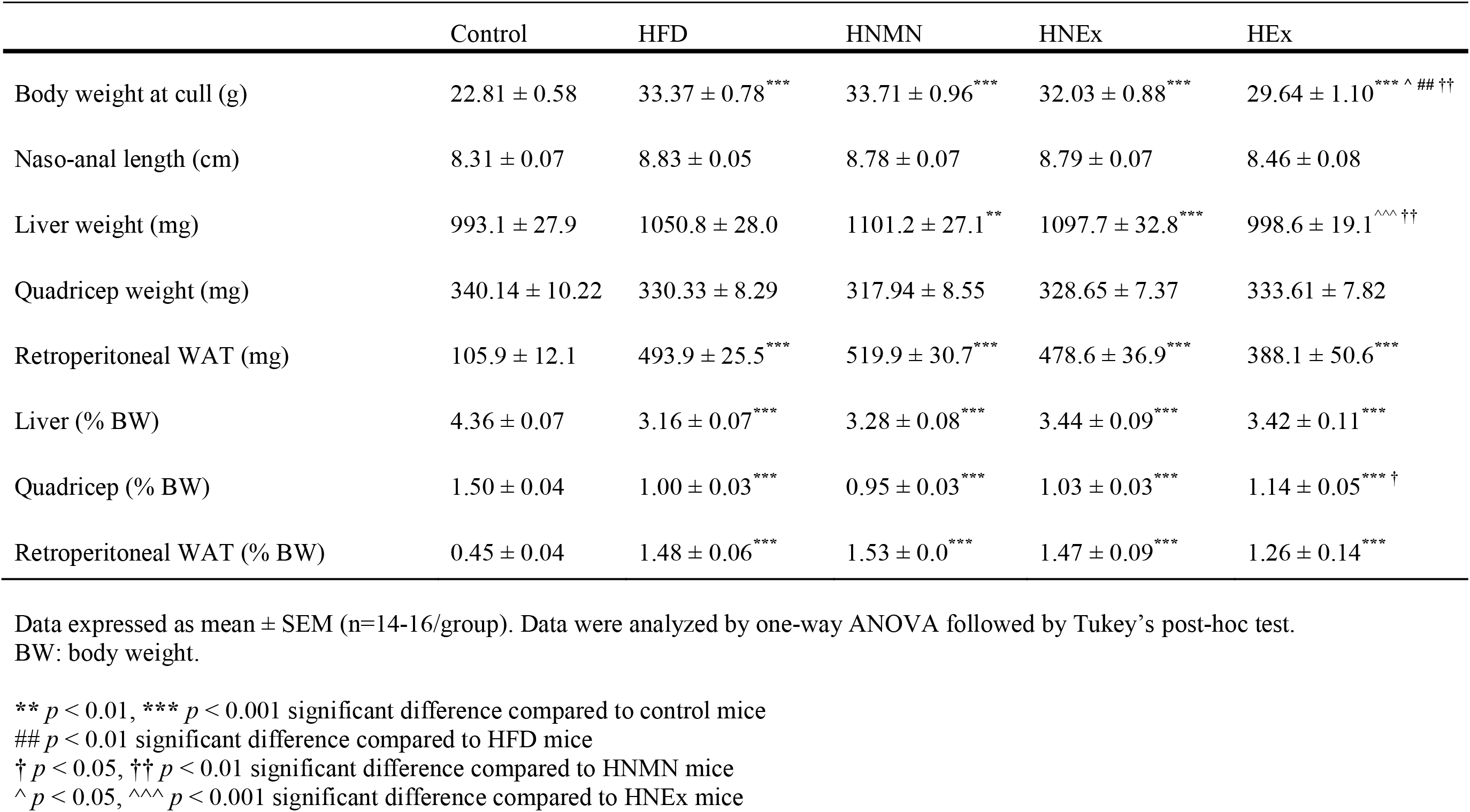
Effects of oral NMN administration and exercise on anthropometric measures in mice consuming HFD.

We observed an increase in net liver mass (**Table 1**) in both the HNMN and HNEx groups compared to control mice. These differences may be partially attributed to an increase in hepatic triglyceride accumulation (**Figure 3A**), however the HFD group also had significantly elevated hepatic triglyceride levels, despite no significant change in liver mass. Interestingly, HEx mice had significantly reduced net liver weight compared to the NMN treated groups and lowered triglyceride content with levels comparable to control mice.

**Figure 3:**
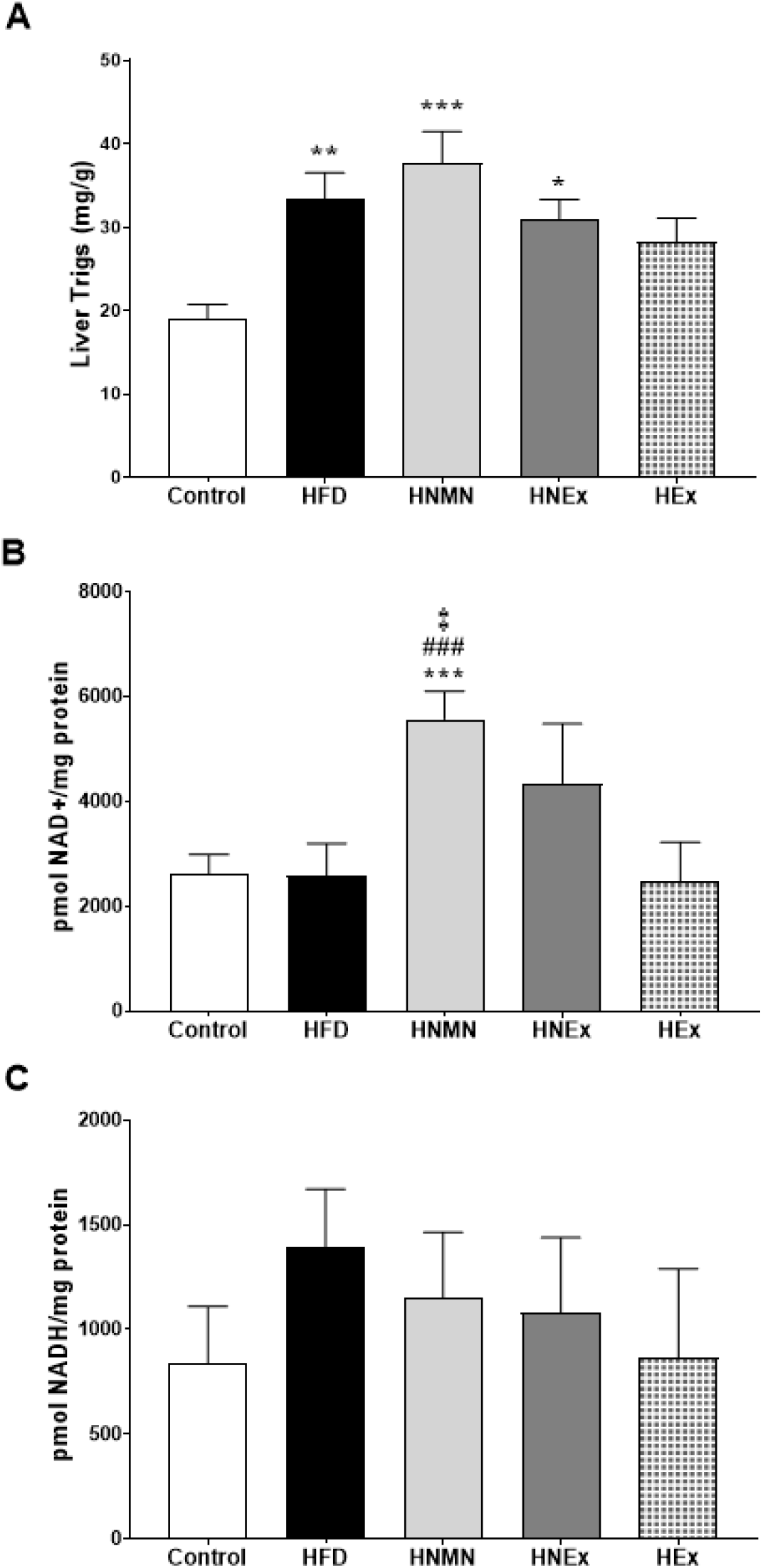
Effects of oral NMN administration and exercise on hepatic triglycerides accumulation (A), NAD^+^ (B) and NADH (C) content. Data expressed as mean ± SEM (n=9-16/group). Data analyzed by one-way ANOVA followed by Tukey’s post-hoc test. *p* < 0.05, ** *p* < 0.01, *** *p* < 0.001 significant difference compared to control mice ### *p* < 0.001 significant difference compared to HFD mice ‡ *p* < 0.05 significant difference compared to HEx mice

When standardized by body weight, no differences were observed across HFD-fed groups (**Table 1**).

### 3.5 Oral Administration of NMN Increased Hepatic NAD^+^ Content

HFD-feeding did not impact hepatic NAD^+^ levels, as measured by an enzyme based NAD cycling assay (**Figure 3B**), consistent with previously published work [30]. Oral NMN administration alone significantly increased hepatic NAD^+^ levels in HNMN compared to chow mice, HFD control and HFD-exercise alone groups. There was also a trend for increased hepatic NAD^+^ levels in the HFD group with combined NMN and exercise treatment. We detected no differences in hepatic NADH levels (**Figure 3C**) among the groups. Metabolomics analyses suggest a trend for increased quadricep NAD^+^ associated metabolites such as increased NAD^+^ content in HNMN mice and NADH in HNEx mice (**Figure S3**).

### 3.6 Effects of oral NMN administration and exercise in skeletal muscle

No differences were observed in mitochondrial DNA content in the quadriceps, in response to HFD suggesting minimal impact on mitochondrial biogenesis (**Figure 4A**). No changes were observed in mice which were only given oral NMN supplementation compared to both non-intervention groups. In contrast, exercise significantly increased quadricep mitochondrial DNA content in both HEx and HNEx mice.

**Figure 4:**
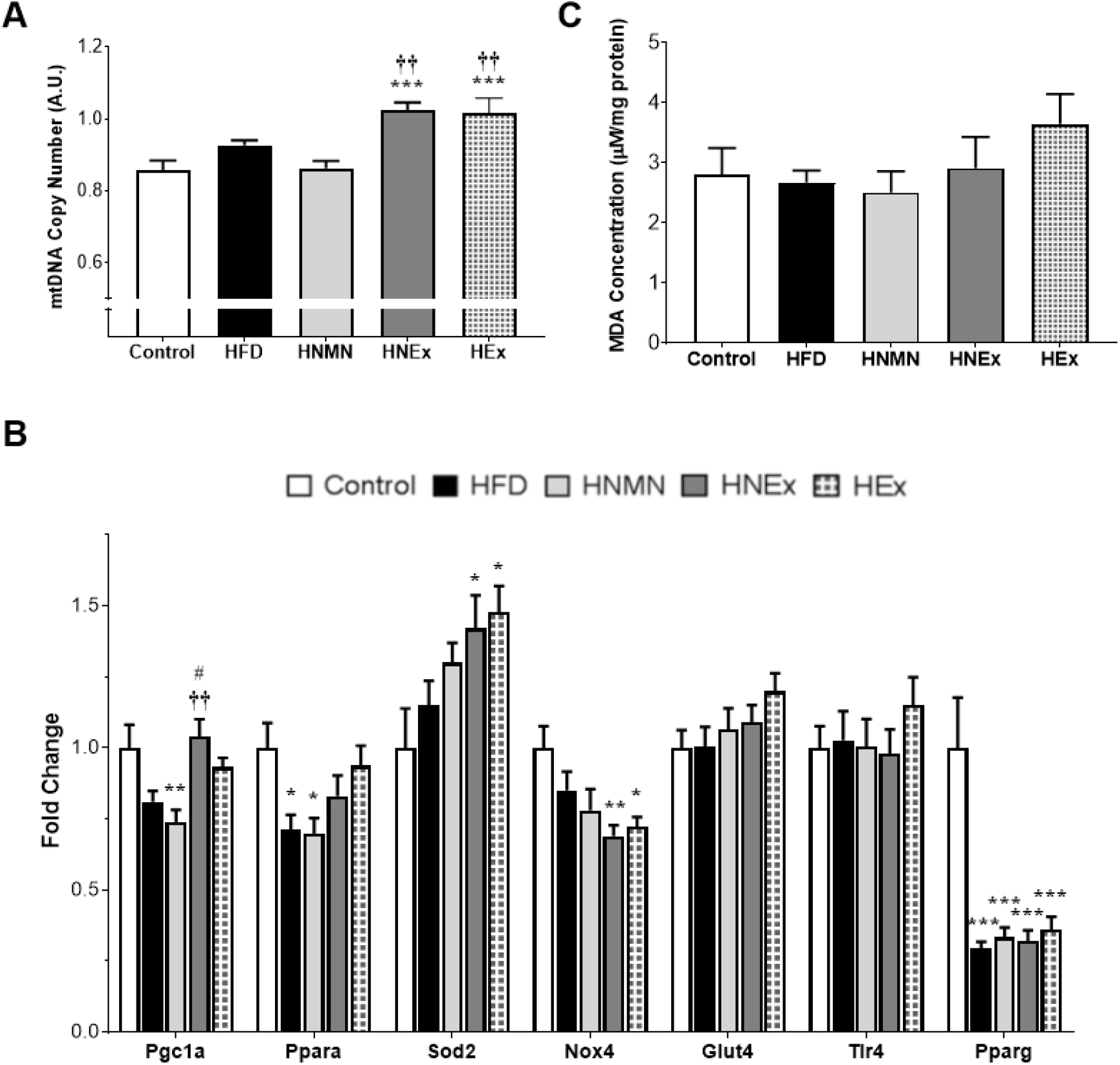
Effects of oral NMN administration and exercise on mitochondrial DNA content (A), gene expression (B) and lipid peroxidation (C) in skeletal muscle. Data expressed as mean ± SEM (n=14-16/group). Data for each gene were normalized to two house-keeper genes and analyzed by one-way ANOVA followed by Tukey’s post-hoc test. \**p* < 0.05, ** *p* < 0.01, *** *p* < 0.001 significant difference compared to control mice # *p* < 0.05 significant difference compared to HFD mice †† *p* < 0.01significant difference compared to HNMN mice

HFD-feeding induced a non-significant decrease in muscle *Pgc1a* expression, which was exacerbated by oral NMN administration (**Figure 4B**). Although exercise alone did not affect *Pgc1a* expression, significant increases were observed in HNEx mice compared to HFD mice and HNMN mice. As mitochondria play a central role in the utilization of fuel for metabolism, we examined *Ppara*, a marker of fat metabolism. As expected, long-term consumption of HFD caused significant reductions in muscle *Ppara* expression compared to standard chow fed mice. No benefits were seen with oral NMN administration alone; HNMN mice had comparable *Ppara* to HFD mice. Exercise was able to ameliorate the reduction caused by HFD-feeding in both HEx and HNEx mice.

Considering *Pgc1a* plays a regulatory role in the expression of antioxidants, *Sod2* and *Nox4* expression were also measured. HFD consumption did not affect the expression of these two genes. However, both groups which received exercise showed significantly increased muscle *Sod2* and decreased *Nox4* expression in comparison to control mice, indicating beneficial effects of exercise on increased antioxidative capacity. Taken together with the increase in *Pgc1a* expression in HNEx mice, the largely increased antioxidative response resembles the dysregulation of redox capacity observed in islet cells, suggesting a possible mechanism for the diminished benefits of exercise in mice also treated with NMN.

Interestingly, analysis of metabolomics data from quadriceps muscle showed a trend for increased coenzyme A (CoA) and components of the TCA cycle, such as succinate and fumarate, in mice which underwent treadmill exercise (**Figure S4**). Mice exposed to exercise alone had a trend for increased malonyl CoA (**Figure S4**), a marker for fatty acid synthesis, and a non-significant increase in quadricep MDA concentration (**Figure 4C**).

## 4. DISCUSSION

For the first time, this study explored the combination of exercise and NMN treatment, both of which have previously shown beneficial effects to replenish the decreased NAD^+^ observed in rodent models of diet-induced obesity. Such approaches are useful to guide current efforts in utilizing NAD-precursor therapies [31, 32]. Here, we demonstrated exercise alone ameliorated impairments from HFD to glucose clearance, as measured by GTT, and to pancreatic beta cell function, as measured by *ex vivo* GSIS. Notably, when exercise was combined with NMN administration, this beneficial effect was dampened. Islet and quadriceps gene expression data showed upregulation of antioxidative genes and a decline in prooxidant genes (*Nox4* and *Gpx4* in islets, *Pgc1a* in quadricep). The increased ratio of antioxidant to prooxidant expression raises the possibility of NMN administration interfering with redox regulation in the context of an exercise intervention in HFD-induced obesity.

Improving metabolic outcomes in obesity by enhancing NAD^+^ availability using precursors is an attractive option to counter the reduced NAD^+^ observed in subcutaneous adipose tissue in human obesity and also in fat pads and livers of rodent obesity models [13, 33]. As the liver plays an important regulatory role in glucose metabolism, studies have explored the hepatic effects of administration of NAD^+^ precursors. Increasing NAD^+^ abundance has been shown to promote mitochondrial biogenesis, consequently causing increased β-oxidation in rodent models [34, 35]. In the present study, despite the increase in hepatic NAD^+^, triglyceride levels were unaffected by oral NMN administration. Potential explanations for this are that in the present study the HFD did not reduce liver NAD^+^ levels, that the dose of NMN was insufficient, or there was too little fat accumulation in the liver, as our previous work demonstrated NMN only reduced liver triglycerides in the most obese mice, following exposure to HFD both *in utero* and post-weaning [36]. Consistent with animal and human studies [37, 38] our data show exercise reduced hepatic triglyceride levels, potentially through increasing beta-oxidation and inhibiting lipogenesis [37, 39, 40]. NMN appeared to dampen this benefit and other measures in this study, explored below.

Glucose intolerance is common in patients with obesity and rodent models of diet-induced obesity [41]. Daily intraperitoneal injections of 500mg/kg NMN improved glucose tolerance, despite no effects on body weight, in rodent models of HFD-induced obesity and diabetes [13, 23]. However, in this study, oral NMN administration alone (400mg/kg) was not sufficient to reduce body weight nor improve glucose tolerance, while treadmill exercise reduced weight gain and improved glucose clearance in HEx mice. Similar benefits of exercise in the context of obesity and diabetes have been demonstrated in rodents and humans [23, 42, 43]. No clear benefits of exercise were observed in mice also given oral NMN, adding to the idea that NMN dulls the effects of exercise.

Given that insulin secretion is critical for maintaining glucose homeostasis, we also investigated effects of exercise and oral NMN administration on pancreatic islets. Insulin secretion in high glucose conditions is impaired in mouse models of obesity [44]. Earlier work by Revollo and colleagues using Nampt-deficient mice [45] established the importance of NAD^+^ in the regulation of β-cell insulin secretion. Subsequent studies have demonstrated the protective effects of NAD^+^ modulation in isolated islets, through pre-incubation with NAD^+^ precursors and enzymes, before functional assessments such as apoptosis and GSIS measures [46, 47]. Here, oral NMN administration in obese mice did not induce any benefits on islet insulin secretion capacity. Others reported a single bolus injection of NMN at 500mg/kg ameliorated the impact of fructose-rich diet feeding on GSIS [48] and improved age-associated decline of GSIS in β-cell specific Sirt-1 overexpressing mice [49]. Thus, in diet-induced obesity, this comparatively low dose of NMN, or route of administration may not have yielded benefits. On the other hand, HEx mice displayed improvements in GSIS response as demonstrated by others [50], but this was reduced in mice also receiving NMN. We examined gene expression data for further clarification. Interestingly, both the metabolic impairment caused by HFD and benefits of exercise were associated with unaltered islet gene-expression under basal conditions. It is possible that metabolism-induced transcriptional changes may occur in glucose-stimulated islets. Nonetheless, under resting conditions, NMN treated groups showed decreased *Nox4* expression. NMN administration also increased expression of *Pgc1a* in islets of HNMN mice and *Gpx4* in HNEx mice, which supports the antioxidative effects of chronic NMN administration. Whilst the mechanism for the benefits of exercise on pancreatic function are still unclear, one proposed mechanism is via differential oxidative and antioxidative responses [51]. Taken together, it is possible that alterations to redox metabolism from NMN intake may impede the benefits of exercise on glucose regulation.

The glucose intolerance and insulin resistance present in obesity can also be attributed to impaired function of skeletal muscle, the organ with greatest glucose uptake and fat oxidation [52]. Here, exercised mice, regardless of NMN administration, had the same compensatory reduction in ambulatory activity as observed in other studies [53, 54], suggesting the potential for skeletal-muscle adaptations to exercise in both groups. Exercise training has been shown to stimulate mitochondrial biogenesis as part of an adaptive mechanism to meet the increased tissue energy requirements and to manage the increased mitochondrial stress instigated through processes such as increased ROS and upregulation of the mitochondrial unfolded protein response [55, 56]. In this study, changes in mtDNA copy number were observed in exercised mice, regardless of NMN administration, which emphasizes the importance of exercise in mitochondrial regulation, and that not all exercise-induced changes were blunted by NMN administration. Our metabolomics data showed a trend for increased malonyl CoA in HEx mice, potentially indicating upregulation of fatty acid synthesis and thus, decreased fatty acid oxidation. A study in endurance athletes demonstrated increased fatty acid synthase mRNA following exercise training, which was associated with *Pgc1a* expression [57].

Interestingly, at a transcriptional level, *Pgc1a* expression was elevated in HNEx mice but did not demonstrate the same increase in malonyl CoA. Upregulation of *Pgc1a* expression may be attributed to regulation by both NAD^+^ dependent SIRT1 deacetylation driven by NMN, and AMPK mediated phosphorylation driven by exercise [21, 58]. In line with our results, a study examining muscle specific over-expression of *Pgc1a* did not find improvements in glucose handling in mice given a western diet, compared to physical exercise [59].

Repeated bouts of contractile activity in skeletal muscle cause stress and remodeling, which triggers the transcription of antioxidant genes as a method of adaptation [60, 61]. MDA levels in the muscle were assessed as a marker for oxidative stress. The trend for an increase we observed in HEx and not HNEx mice provide further support for the antioxidative effects of NMN administration which, in HNEx mice, may disrupt the post-exercise surge in ROS which is necessary for health benefits.

Studies using conventional antioxidants have shown attenuation of gene expression changes induced by exercise in skeletal muscle [62, 63]. It has been well established that NAD^+^ is an essential cofactor for antioxidant defense, stimulating SIRT activity and ultimately allowing for cytosolic production of NADPH from NADP^+^. The cycling of NADPH to NADP^+^ is critical for the recycling of glutathione and thioredoxin from their oxidized states [64]. Thus, we considered the possibility of oral NMN administration increasing cellular NAD^+^ levels and consequently antioxidant defense mechanisms. The trend seen in the quadriceps metabolomics data for increased NADPH in mice given oral NMN may also support the hypothesis that NMN boosts post-exercise antioxidant function. This may not occur at the transcriptional level as we found that exercise-induced gene changes in *Nox4* and *Sod2* were similar in the quadriceps of HEx and HNEx mice. Again, our data do not exclude alteration of the acute post-exercise transcriptional response. An aim of future work would be to test this immediate post-exercise window to see if NMN alters oxidative stress ultimately leading to reduction in metabolic improvements such as in glucose tolerance [65, 66].

Another explanation for the abrogation of exercise benefits from NMN treatment could be altered ROS signaling. ROS plays an important signaling role in processes such as insulin secretion in islets and sensitivity in skeletal muscle, and treatment with antioxidants abolishes the metabolic benefits of exercise [60, 61]. It is possible that the increased expression of antioxidant enzymes observed during NMN treatment impaired ROS signaling, blocking metabolic or mitochondrial benefits from exercise.

It is notable that in our model, neither muscle nor liver NAD^+^ levels were reduced in HFD fed mice which may explain why supplementing with the NAD^+^ precursor NMN had no positive impact on mice which were exercised. Massive over-expression of the NAD biosynthetic enzyme NAMPT leads to only moderate changes in NAD levels [67, 68], and it has been argued that NAD levels are tightly regulated such that increased NAD synthesis results in a commensurate increase in NAD breakdown [69]. Exogenous NMN treatment in animals where NAD^+^ levels are not impaired may result in increased NAD^+^ breakdown to maintain NAD^+^ homeostasis, and while treatment with NAD^+^ precursors are increasingly seen as broadly beneficial, this may be less effective in situations where NAD^+^ levels are replete.

## 5. CONCLUSIONS

In conclusion, our data show exercise alone ameliorated metabolic impairments caused by HFD. Whilst oral administration of NMN alone increased NAD^+^ levels, the absence of improved metabolic outcomes could be due to our HFD not inducing a deficiency in NAD^+^ compared to control. However, our data suggest that NMN administration at 400mg/kg in drinking water could drive an antioxidative response which may have mitigated the transient increase in ROS after exercise, which is crucial for exercise-induced health benefits such as improved insulin secretion in response to glucose stimulation and metabolic adaptation. These results argue against the idea that combining NMN-supplementation and exercise would have additive benefits in the treatment of obesity, and warrant caution in the use of NAD+ precursors in situations where NAD+ is already replete.

## Abbreviations

HFD: high fat diet
ROS: reactive oxygen species
ATP: adenosine triphosphate
PARP: poly (ADP-ribose) polymerase
SIRT: sirtuins
NAD^+^: nicotinamide adenine dinucleotide
NMN: nicotinamide mononucleotide

## AUTHOR CONTRIBUTIONS

M.J.M. and N.A.Y. designed the study. J.Y., LJ.K., LE.Q. and N.A.Y. conducted experiments. J.Y., D.R.L., M.J.M. and N.A.Y. interpreted results and wrote the manuscript. All authors contributed to editing the manuscript and approved the manuscript for publication.

## ACKNOWLEDGEMENTS

This study was supported by the Rebecca Cooper Foundation grant to N.A.Y., National Health and Medical Research Council (NHMRC) Australia project funding to M.J.M., NHMRC Career Development Fellowship (APP1127821) to L.E.W., and J.Y. is supported by the Research Training Program, UNSW Sydney.

## DECLARATION OF INTEREST

L.E.W. is an inventor on patents licensed to Metro Biotech NSW and to Jumpstart Fertility, and has received sponsored research funding from both companies. He is an advisor and shareholder in EdenRoc Sciences (Metro Biotech NSW, Metro Biotech, Liberty Biosecurity), Life Biosciences LLC and its daughter companies (Jumpstart Fertility, Continuum Biosciences, Senolytic Therapeutics, Selphagy, Animal Biosciences, Iduna) and in Intravital Pty Ltd. L.E.W. provides consulting work for Life Biosciences.

