## Supplementary Figures for "EXERCISE-INDUCED BENEFITS ON GLUCOSE HANDLING IN A MODEL OF DIET-INDUCED OBESITY ARE REDUCED BY CONCURRENT NICOTINAMIDE MONONUCLEOTIDE"

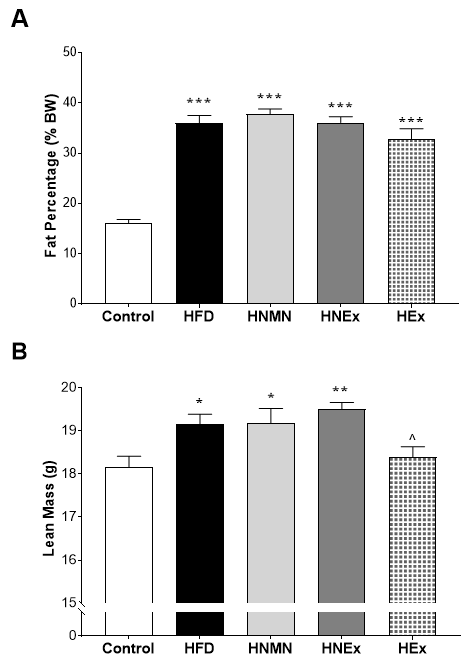

### SUPPLEMENTARY FIGURES

**Supplementary Figure 1: Effects of oral NMN administration and exercise on body fat percentage (A) and lean mass (B).** Data analysed by one-way ANOVA followed by Tukey’s post-hoc test.

***** *p* < 0.05, ****** *p* < 0.01, ******* *p* < 0.001 significant difference compared to control mice

^ *p* < 0.05 significant difference compared to HNEx mice

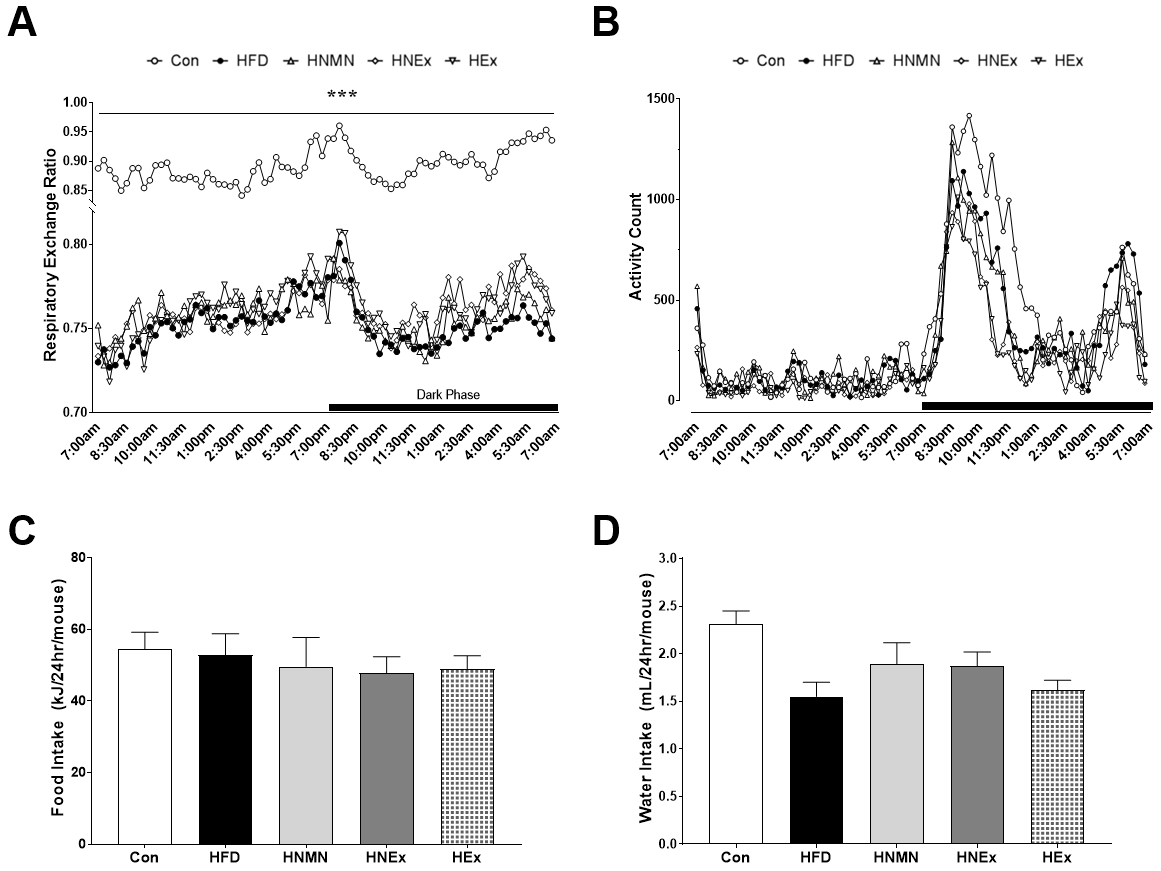

**Supplementary Figure 2: Calorimetry measures from CLAMS showing respiratory exchange ratio (RER; A) and ambulatory activity (B), as well as individual 24-hour food (C) and water (D) intake**. Data expressed as mean ± SEM (n=12/group). RER and activity count analysed by repeated measures ANOVA; individual food and water intake analysed by one-way ANOVA followed by Tukey’s post-hoc test.

******* *p* < 0.001 significant difference compared to control mice

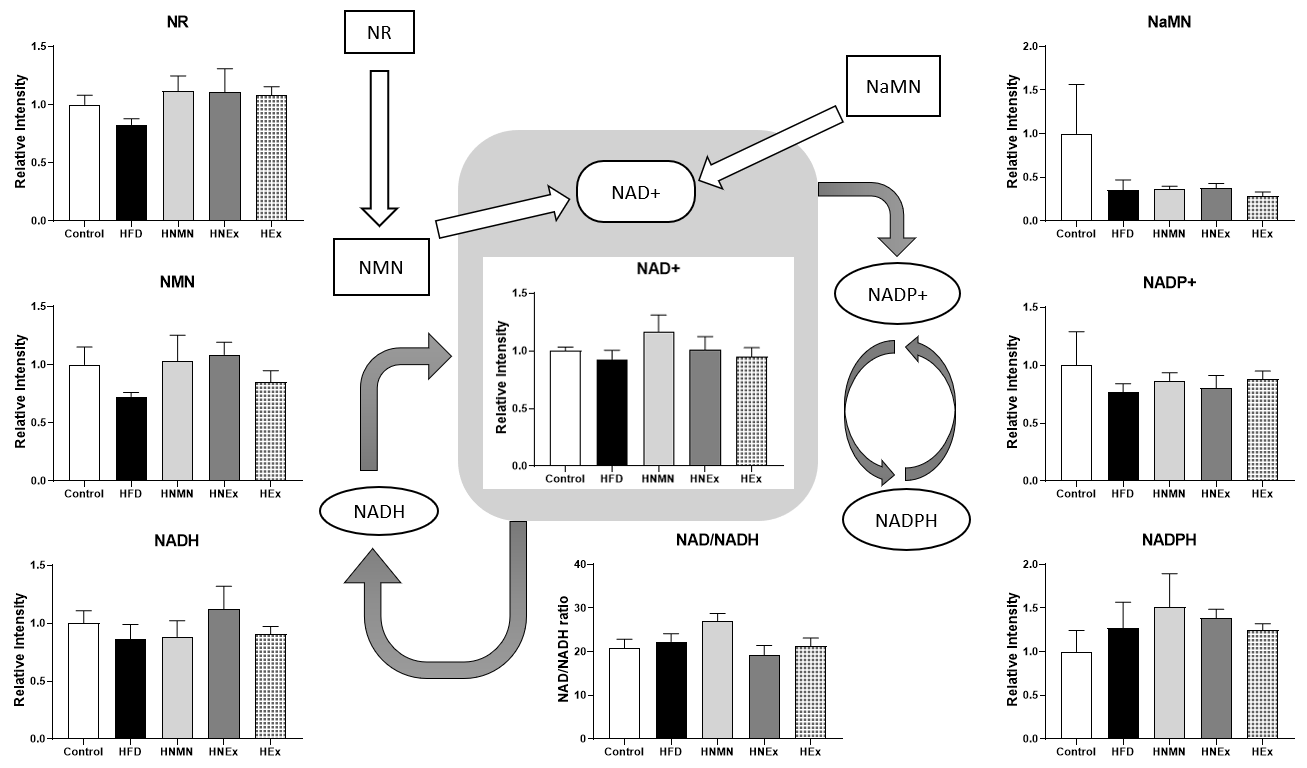

**Supplementary Figure 3: NAD^+^ metabolites in quadriceps analysed by LC-MS/MS and a simplified schematic of NAD^+^ pathway.** NAD^+^ precursors shown in boxes, and cofactors central to metabolism which utilise NAD^+^ shown in ovals. Data expressed as mean ± SEM (n=3-4/group). Data for each metabolite were normalized to internal standard and tissue weight, then analysed by one-way ANOVA followed by Tukey’s post-hoc test.

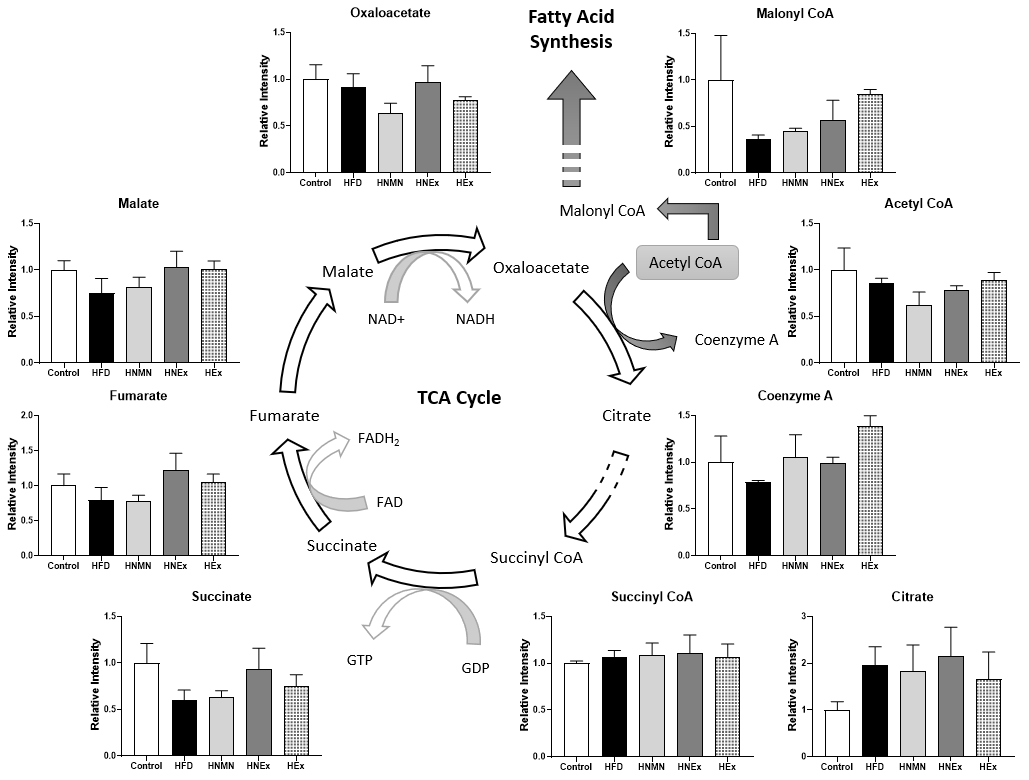

**Supplementary Figure 4: Metabolites involved in the TCA cycle and fatty acid synthesis, in quadriceps, analysed by LC-MS/MS against a simplified schematic of the TCA cycle.** Acetyl CoA reacts with oxaloacetate, as the first step of the TCA cycle, to create citrate and produce coenzyme A. Acetyl CoA can also be utilised for fatty acid synthesis via malonyl CoA. Data expressed as mean ± SEM (n=3-4/group). Data for each metabolite were normalized to internal standard and tissue weight, then analysed by one-way ANOVA followed by Tukey’s post-hoc test.
